# *Candida albicans* white and opaque cells exhibit distinct spectra of organ colonization in mouse models of infection

**DOI:** 10.1101/461293

**Authors:** Julie Takagi, Sheena D. Singh-Babak, Matthew B. Lohse, Chiraj K. Dalal, Alexander D. Johnson

## Abstract

*Candida albicans*, a species of fungi, can thrive in diverse niches of its mammalian hosts; it is a normal resident of the GI tract and mucosal surfaces but it can also enter the bloodstream and colonize internal organs causing serious disease. The ability of *C. albicans* to thrive in these different host environments has been attributed, at least in part, to its ability to assume different morphological forms. In this work, we examine one such morphological change known as white-opaque switching. White cells are the default state of *C. albicans*, and most animal studies have been carried out exclusively with white cells. Here, we compared the proliferation of white and opaque cells in two murine models of infection and also monitored, using specially constructed strains, switching between the two states in the host. We found that white cells outcompeted opaque cells in many niches; however, we show for the first time that in some organs (specifically, the heart and spleen), opaque cells competed favorably with white cells and, when injected on their own, could colonize these organs. In environments where the introduced white cells outcompeted the introduced opaque cells, we observed high rates of opaque-to-white switching. We did not observe white-to-opaque switching in any of the niches we examined.

## Introduction

*Candida albicans* is a source of serious fungal infections in the United States; it is the fourth most commonly cultured microbe from blood (Pfaller and Diekema, 2007). Unlike other major fungal pathogens, *C. albicans* is also a part of the normal human microbiome, particularly in the gastrointestinal tract, skin, oral cavity, and other mucosal surfaces (Odds, 1988). As a pathogen, *C. albicans* poses health risks, particularly for people whose immune system is suppressed, who have surgeries, implanted medical devices, or who have been treated with long courses of antibiotics. The attributable mortality from *C. albicans* bloodstream infections in adults is at least 15% and has been reported to be as high as 40% (Zaoutis et al., 2005).

While some human fungal pathogens exist primarily as budding yeast cells (for example, *Cryptococcus neoformans* (Buchanan, 1998)) or filamentous hyphal structures (for example, *Aspergillus* spp. (Baker and Bennett, 2007)), *C. albicans* alternates between these and other morphologies, often in response to specific environmental cues. In this paper, we explore the switch *C. albicans* makes from its normal, round-to-oval yeast cell morphology (white) to an elongated, mating competent cell type termed opaque (for reviews see Lohse and Johnson, 2009; Soll et al., 2009). White-opaque switching is unusual in that it is a heritable change, one that occurs without a change in the DNA sequence of the genome. In standard laboratory media, switching occurs rarely, approximately once every 10^4^ cell divisions (Slutsky et al., 1987; Soll et al., 1993); therefore, each cell type is accurately maintained across many cycles of cell division. Approximately one-sixth of all genes (of ~6000 total) are differentially regulated (at least two-fold) between the two cell types (Lan et al., 2002; Tuch et al., 2010). These expression differences involve many types of genes, including those responsible for the ability of opaque cells (but not white cells) to mate, consistent with the observation that opaque cells are specialized for mating (Miller and Johnson, 2002; Tsong et al., 2003). The complete set of differentially regulated genes also encodes a variety of metabolic enzymes, suggesting that white and opaque cells are each ‘metabolically specialized’ to thrive in specific niches (reviewed in Huang, 2012; Lohse and Johnson, 2009; Morschhäuser, 2010; Soll, 2014, 1992, 2004). Supporting this hypothesis, environmental factors such as temperature, carbon source, and oxygen can significantly alter the switching rate between cell-types (Ene et al., 2016; Huang et al., 2009, 2010; Rikkerink et al., 1988). Another difference between white and opaque cells lies in their recognition by macrophages and neutrophils, indicating that the white-opaque switch may play a role in evading the host innate immune system response and in colonizing internal environments (Geiger et al., 2004; Kolotila and Diamond, 1990; Kvaal et al., 1999, 1997; Lohse and Johnson, 2008; Sasse et al., 2013). Taken as a group, these observations point to the hypothesis that white-opaque switching evolved during *C. albicans*’ long association with its warm-blooded host and plays a crucial role in the pathogen-host relationship. Consistent with this hypothesis, deletion of the master regulator of white-opaque switching (Wor1) impairs the ability of *C. albicans* to survive in a mouse gastrointestinal model of colonization (Pande et al., 2013). Although not all clinically isolated strains of *C. albicans* readily undergo white-opaque switching, many do so. Even “non-switching” clinical strains can be converted to switching strains by simple mutation, indicating that the complex mechanism underlying switching and the specification of the two cell types are deeply conserved traits. Also consistent with this hypothesis is the observation that two pathogenic fungi closely related to *C. albicans* (*Candida dubliniensis* and *Candida tropicalis*) also undergo white-opaque switching (Porman et al., 2013; Pujol et al., 2004).

To systematically investigate the behavior of white and opaque cells *in vivo*, genetically identical strains of white and opaque cells were tagged with spectrally distinct fluorophores and co-inoculated into rodent models of infection and colonization. Historically, these animal models have been optimized for white cells, and in this study, we varied the dosage and timing to better capture the behavior of opaque cells. We found, using variations of the tail-vein injection model (Odds et al., 2000), that both white and opaque cells are capable of disseminating from the bloodstream into multiple organs; in some organs, white cells significantly outcompete opaque cells, but in others, they are roughly equivalent. In gut models of colonization (Rosenbach et al., 2010), with and without antibiotic treatment, long-term colonization by cells introduced as opaque cells depends on their switching to white cells.

## Results

### Fluorescent markers enable unbiased comparison of white and opaque cells *in vivo*

In order to develop a quantitative, reproducible approach to observe the relative colonization of white and opaque cells *in vivo*, we constructed fluorescently tagged white and opaque strains of *C. albicans*. These strains were matched genetically, except for a single locus where the Tef2 protein was C-terminally tagged with either GFP or mCherry, two spectrally distinct fluorophores. We employed experiments with a plate reader to validate that tagging white and opaque cells with fluorophores did not affect their growth (Figure S1, Materials and Methods, Dataset 1). These strains were then introduced together (in 50:50 mixes) to mice via tail-vein injection or oral gavage; after the mice were sacrificed, *C. albicans* cells were harvested and immediately plated at 25°C. After seven days of growth, we scored colonies (based on their colony morphology) as white or opaque to determine the state of each cell at the point the mice were sacrificed. (At 25°C on plates, very little switching occurs, so the appearance of a given colony reflected the identity of the single cell that produced it.) Using a fluorescent stereomicroscope, we also determined whether each colony expressed GFP or mCherry. In this way, we were able to determine whether a given cell was injected as a white or opaque cell and whether it remained as the same cell type or underwent a switch to the other cell type. We validated this approach by plating mixtures of white and opaque cells with known ratios; we were able to recover the expected ratio of colonies on plates (Figure S2, Materials and Methods, Dataset 2). Hence, this approach offers several advantages when compared with previous studies: 1) it does not require the use of auxotrophies or drug resistance markers that have been shown to affect *C. albicans* behavior *in vivo* (Bain et al., 2001; Lay et al., 1998), 2) it does not require the use of strains that are artificially “locked” in one cell-type or another, for example by deleting or overexpressing the master regulator Wor1 (Huang et al., 2006; Pande et al., 2013; Zordan et al., 2006), and 3) it does not require any downstream processing, such as qPCR, which can introduce errors through amplification. Finally, because white and opaque cells were injected together, this approach enabled us to directly compare colonization, proliferation, and switching of white and opaque cells *in vivo*.

To test whether the fluorophores introduced any bias *in vivo*, the tail veins of four Balb/C mice (18-20g) were injected with 2.2*10^6^ white cells, half of which were tagged with GFP and the other half with mCherry. The mice were sacrificed after 24 hours and the fungal burden in five organs (the kidney, liver, spleen, heart, and brain) was analyzed (Figure 1a). Across all five organs, the distributions of mCherry and GFP-tagged colonies were not significantly different (p-values all >0.05, Wilcoxon matched-pairs signed rank test, Figure 1, Dataset 3), indicating that the fluorophores themselves introduce little or no bias *in vivo*.

**Figure 1.**
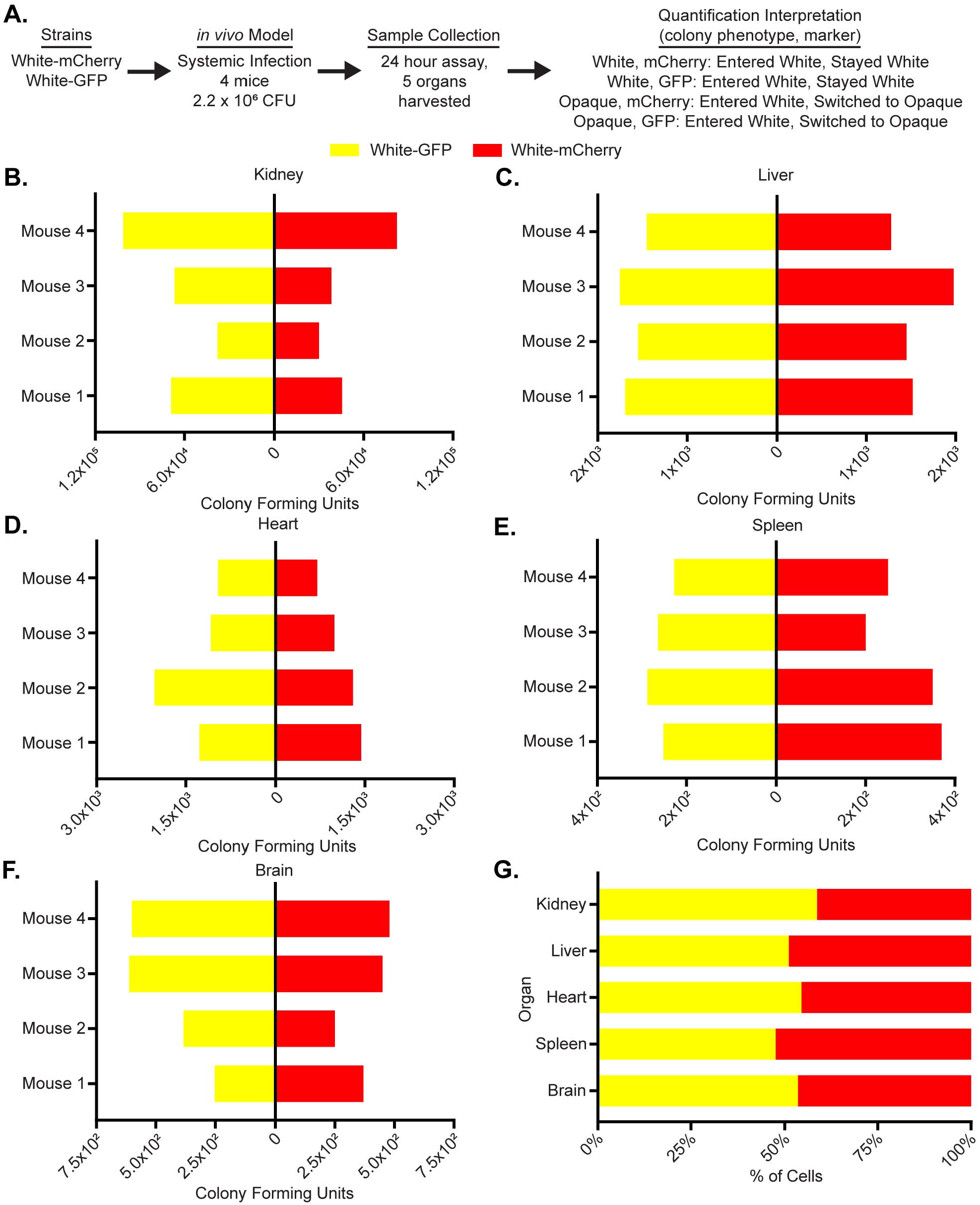
The use of fluorescent reporters does not influence *C. albicans* activity *in vivo.* (a) Using a flowchart, the experimental setup, cell type, and potential fluorescence phenotypes for each strain are tabulated. In this case, white cells expressing GFP and mCherry were co-injected into the tail-veins of 4 mice. Five organs, the kidney, liver, heart, spleen and brain, were processed to measure white and opaque cell colonization as well as white-opaque switching. The mechanistic interpretation of each phenotype; in other words, whether or not it indicates cell-type switching, is also indicated. The colony-forming units of white cells expressing GFP (yellow) and the colony-forming units of white cells expressing mCherry (red) are plotted for each mouse for the (b) kidney, (c) liver, (d) heart, (e) spleen and (f) brain. (g) The mean percentage (from n=4 mice) of total cells that are white cells expressing GFP (yellow) and white cells expressing mCherry (red) are plotted per organ as a horizontal bar graph. The raw data for this experiment is available in Dataset 3.

### Opaque cells can disseminate from the bloodstream into many organs

To compare the relative colonization of white and opaque cells in a systemic model of infection, white-mCherry and opaque-GFP cells were co-injected (at a 50:50 ratio) into the tail vein of four mice each at two inocula: 8*10^5^ and 1.6*10^6^ cells (Figure 2a). After 24 hours, the mice were euthanized and their organs harvested and processed. Both white and opaque cells were present in all five organs (as indicated by the colony counts) and were at a higher density in the mice with the higher inoculum (Figure 2, Dataset 4). A comparison of the white and opaque cells across the five organs showed different distributions depending on the organ (Figure 2, Dataset 4. The most pronounced difference was observed in the kidney, where white cells greatly outnumbered opaque cells in every mouse examined (p-value = 0.0078, Wilcoxon matched-pairs signed rank test, Figure 2b, Dataset 4). Of the white cells recovered from the kidney, 70% had been injected as white cells; the remaining 30% had been initially injected as opaque cells but had switched to white cells. The situation in the heart and spleen was different; here, opaque cells outnumbered white cells (p-value 0.0156, Wilcoxon matched-pairs signed rank test, Figure 2d, 2e Dataset 4). In the heart, virtually none of the cells had switched in either direction; in the spleen, a small fraction of the white cells recovered had been injected as opaque cells. Finally, in the liver and brain, both white and opaque cells were recovered with a slight preference for white cells (Figure 2c, 2f). Because of significant mouse-to-mouse variation (particularly in the liver), we do not feel confident that this preference is biologically meaningful even though it is statistically significant (p-values =0.0078, Wilcoxon matched-pairs signed rank test, Figure 2c, 2f, Dataset 4). Rather, we view the colonization of white and opaque cells in these organs as roughly equivalent. We do note that, of the white cells recovered from the brain, a significant fraction had been initially injected as opaque cells.

**Figure 2.**
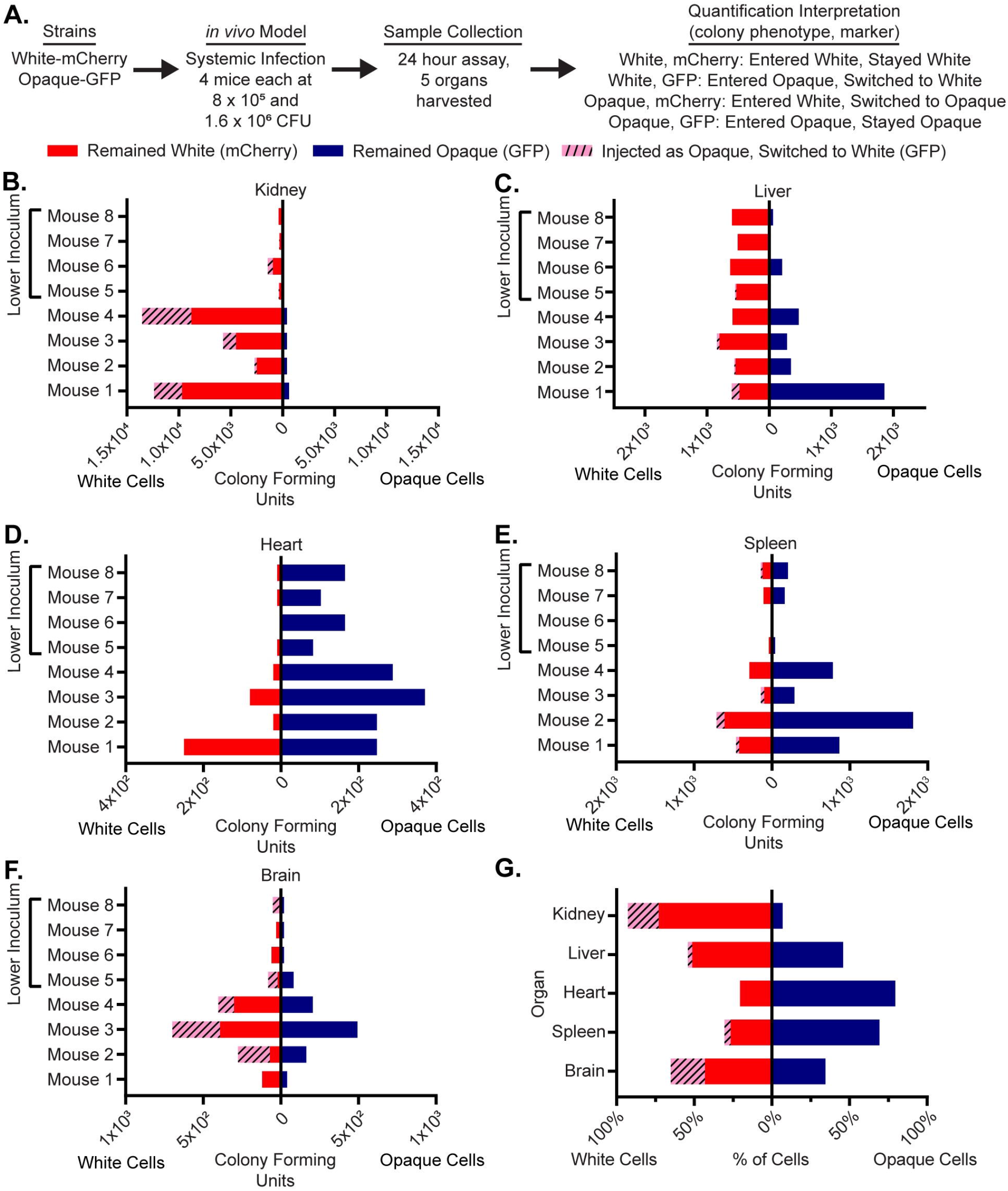
Opaque cells can colonize multiple organs but white cells are preferred in the kidney. (a) Using a flowchart, the experimental setup, cell type, and potential fluorescence phenotypes for each strain are tabulated. In this case, white cells expressing mCherry and opaque cells expressing GFP were co-injected into the tail-veins of 8 mice. Five organs, the kidney, liver, heart, spleen and brain, were processed to measure white and opaque cell colonization as well as white-opaque switching. The mechanistic interpretation of each phenotype; in other words, whether or not it indicates cell-type switching, is also indicated. The colony-forming units of cells that remained white (i.e. white cells expressing mCherry, red), that remained opaque (i.e. opaque cells expressing GFP, blue), and that switched from opaque-to-white (i.e. white cells expressing GFP, dashed pink) are plotted for each mouse for the (b) kidney, (c) liver, (d) heart, (e) spleen and (f) brain. The left side of each horizontal bar graph refers to cells that were white at the end of the experiment while the right side of each horizontal bar graph refers to cells that were opaque at the end of the experiment. The four mice that received the lower inoculums are indicated in each panel. (g) The mean percentage of total cells that remained white (i.e. white cells expressing mCherry, red), that remained opaque (i.e. opaque cells expressing GFP, blue), and that switched from opaque-to-white (i.e. white cells expressing GFP, dashed pink) are plotted per organ as a horizontal bar graph. The left side of the horizontal bar graph refers to cells that were white at the end of the experiment while the right side refers to cells that were opaque at the end of the experiment. The raw data for this experiment is available in Dataset 4.

As described above, our observations document a significant level of opaque-to-white switching that occurred *in vivo*. We do not know when, during the course of the infection, this switching occurred; however, we suspect it happened after the injected cells entered the different organs. Had the opaque cells switched in the bloodstream immediately following injection, we would have expected to observe similar opaque-to-white switching patterns across all organs tested, and this was not the case. We also note that we did not observe any opaque-mCherry colonies (in any organ, in any mouse, and at any inoculum, >1000 colony-forming units visually inspected), indicating that stable switching from white cells to opaque cells must be rare, below the detection limit of the mouse models we examined.

Finally, our data show that in this experiment (Figure 2), mice were inoculated with more white than opaque cells (2:1 white:opaque ratio, see Dataset 4 for raw data). We find that this did not affect strain competition *in vivo* since (1) we found instances (for example, the heart and spleen) where opaque cells grew well, even when they were underrepresented in the inoculum, and (2) this experiment was conducted at two inoculums (Figure 2, Dataset 4) and the trends from both inoculums were the same suggesting that changing the absolute number of either white or opaque cells had no bearing on the relative colonization of different organs.

To test whether the colonization by opaque cells was dependent on co-injection with white cells, we injected the tail veins of four additional mice with a 50:50 mixture of opaque-mCherry and opaque-GFP inoculum (1.6*10^6^ cells) and measured colonization in the kidney, spleen, liver, brain, and heart (Figure S3, Dataset 4). We observed opaque cell colonization of all five organs (Figure S3, Dataset 4), indicating that a large white cell population was not needed to “aid” the opaque cells in colonization.

Taken together, these results show that opaque cells are capable of efficiently colonizing and proliferating in numerous niches *in vivo*, and in some cases (heart and spleen) somewhat better than white cells. However, opaque cells are severely outcompeted by white cells in the kidney, the organ that is typically used to assess systemic fungal infection in murine models.

### *C. albicans* cells that persist for long times during systemic infection are predominantly white

In the previous experiments, we used a relatively large inocula of *C. albicans* and sacrificed the mice after 24 hours. In the next set of experiments, we compared white and opaque cells in a longer-term infection model initiated with a lower inoculum. This model is the standard systemic model of fungal infection (Noble et al., 2010; Odds et al., 2000; Perez et al., 2013). Seven mice were infected with 5*10^5^ cells containing an approximately equal number of white-mCherry and opaque-GFP cells. Mice were sacrificed upon signs of sickness (occurring between 7 and 14 days after injection) and fungal burden was assessed across the five organs as described above (Figure 3a). In this experiment, *C. albicans* cells were recovered reproducibly only from the kidney; either the *Candida* cells were cleared from the other organs or the inoculum was sufficiently low that *Candida* did not efficiently disseminate to the other organs. We also attempted to measure white and opaque cells in the bloodstream during this experiment but were unable to recover a significant number of colony-forming units, consistent with what has previously been observed for the tail-vein injection model (van Deventer et al., 1995; Lionakis et al., 2011).

**Figure 3.**
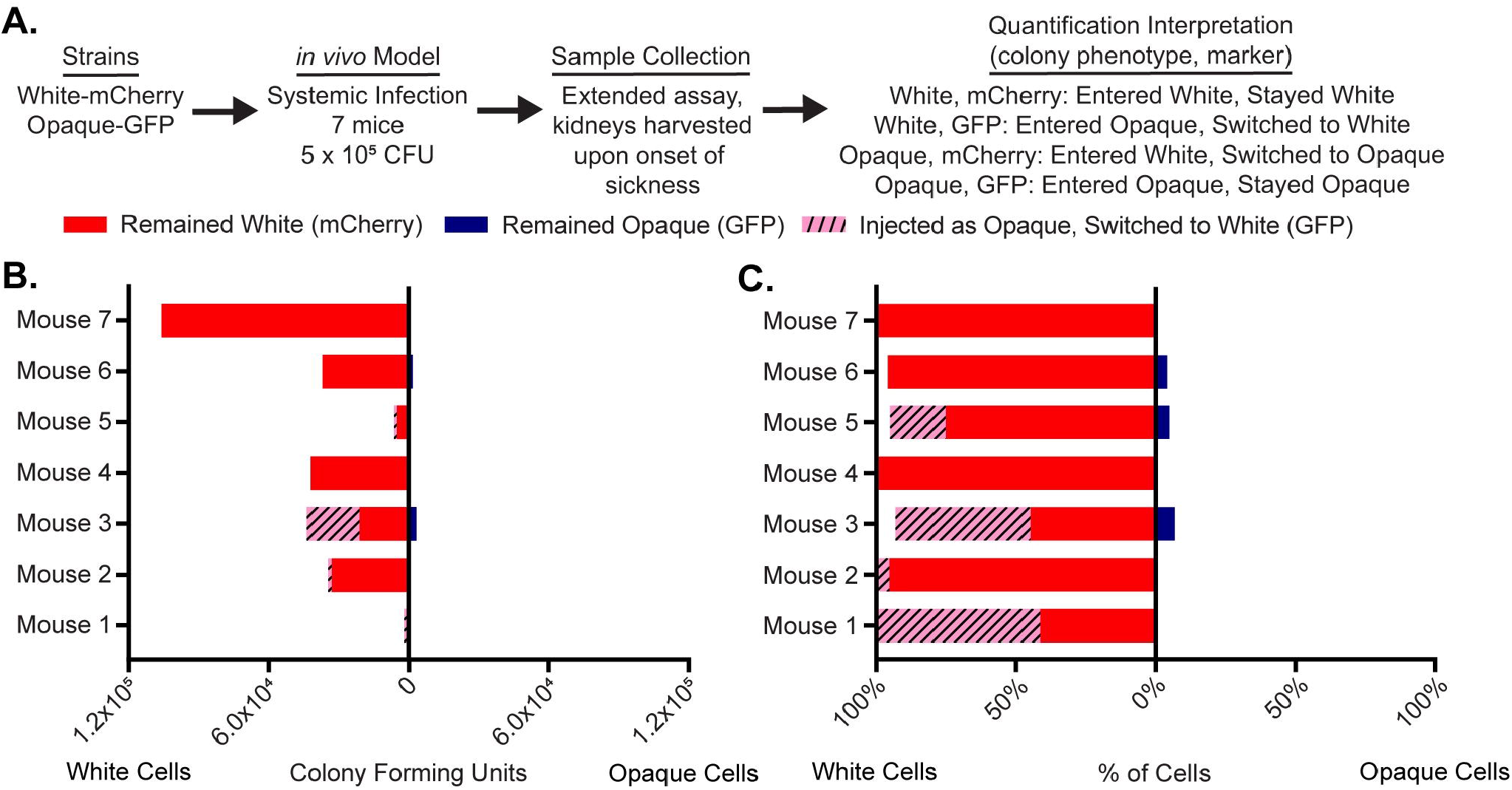
White cells are preferred in the kidney in long-term infections. (a) Using a flowchart, the experimental setup, cell type, and potential fluorescence phenotypes for each strain are tabulated. In this case, white cells expressing mCherry and opaque cells expressing GFP were co-injected into the tail-veins of 7 mice. Once the mice became ill, the kidney was processed to measure white and opaque cell colonization as well as white-opaque switching. The mechanistic interpretation of each phenotype; in other words, whether or not it indicates cell-type switching, is also indicated. (b) The number of colony-forming units of cells that remained white (i.e. white cells expressing mCherry, red), that remained opaque (i.e. opaque cells expressing GFP, blue), and that switched from opaque-to-white (i.e. white cells expressing GFP, dashed pink) are plotted per mouse as a horizontal bar graph. The left side of the horizontal bar graph refers to cells that were white at the end of the experiment while the right side refers to cells that were opaque at the end of the experiment. (c) The percentage of total cells that remained white (i.e. white cells expressing mCherry, red), that remained opaque (i.e. opaque cells expressing GFP, blue), and that switched from opaque-to-white (i.e. white cells expressing GFP, dashed pink) are plotted for each mouse as a horizontal bar graph. The left side of the horizontal bar graph refers to cells that were white at the end of the experiment while the right side refers to cells that were opaque at the end of the experiment. The raw data for this experiment is available in Dataset 5.

Consistent with the previous experiments, cells recovered from the kidney were overwhelmingly white. In several mice, a significant portion of the white cells isolated from the kidney had been initially injected as opaque cells indicating significant opaque-to-white switching *in vivo* (0-58% of cells, depending on the mouse, Figure 3b, 3c, Dataset 5). We also performed an experiment with an inoculum comprised only of opaque cells, half of which were marked with GFP and half of which were marked with mCherry (6.2 * 10^5^ cells). Both populations were recovered from the kidney, and most recovered cells had switched from opaque-to-white (Figure S4). We did observe an unanticipated outcome of this experiment. Rather than an equal ratio of the two strains (as was present in the inoculum), the majority of cells in each individual mouse expressed either GFP or mCherry. We speculate that this observation reflects the stochastic nature of white-opaque switching: there would be a strong bias towards colonization by whichever fluorescent strain switched first.

Consistent with this interpretation is an additional control experiment: mice injected with an inoculum containing an equal number of white-GFP and white-mCherry cells (5*10^5^ cells total) resulted in approximately equal numbers of GFP and mCherry-expressing white cells in the kidneys of three of four mice (Figure S5).

### Opaque cells switch to white cells in an intestinal colonization model

To investigate how white and opaque cells compare in GI tract colonization, we utilized the standard intestinal colonization model (Pande et al., 2013; Perez et al., 2013; Rosenbach et al., 2010). Here, we introduced an inoculum comprised of an equal ratio of white and opaque cells by oral gavage (5*10^7^ cells total per mouse, n=8 mice, Figure 4a). Colonization was monitored by collecting fecal pellets at various time points post-inoculation. We observed that nearly all the introduced opaque cells had switched to white cells within 24 hours and remained white until the mice were sacrificed after 17 days (Figure 4b). All cells injected as white cells remained as white cells throughout the experiment; we did not observe any instance of white-to-opaque switching (>3500 colony-forming units observed). We note that these results mirror, at least superficially, those observed in the kidney following tail-vein injection infections described above. Moreover, they are consistent with results from the oropharyngeal candidiasis model, where introduced opaque cells were either cleared from the oropharynx or switched rapidly to white cells (Solis et al., 2018). Because the observation that opaque cells do not compete well with white cells in the GI tract is entirely consistent with what has been previously published (Böhm et al., 2017), we conclude that tagging white and opaque cells with fluorophores did not introduce any bias that affects strain competition in the GI tract.

**Figure 4:**
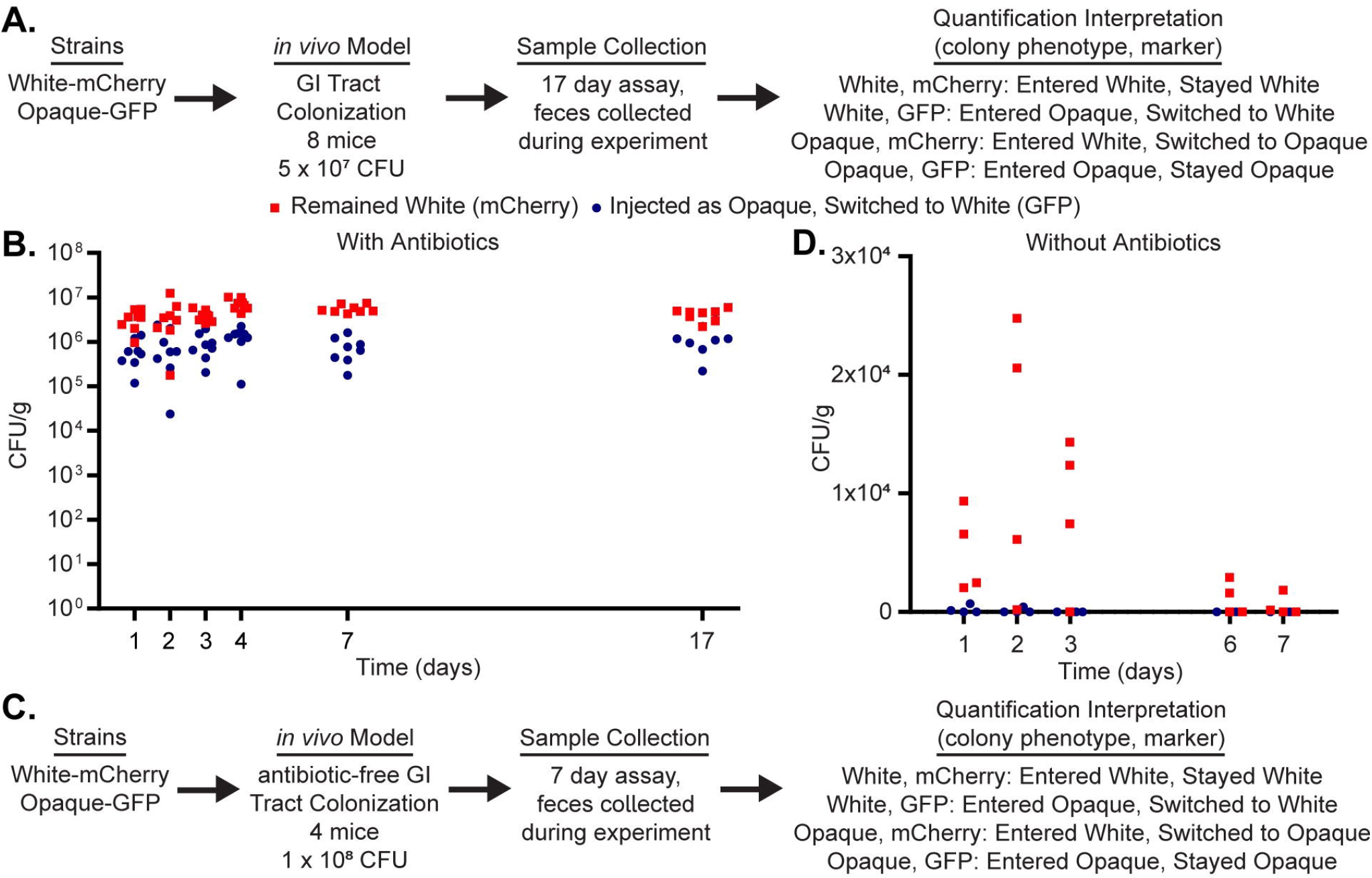
White cells are preferred in an intestinal colonization model of infection. (a) Using a flowchart, the experimental setup, cell type, and potential fluorescence phenotypes for each strain are tabulated. In this case, white cells expressing mCherry and opaque cells expressing GFP were inoculated into 8 mice via oral gavage. Fecal pellets were processed to measure white and opaque cell colonization as well as white-opaque switching. The mechanistic interpretation of each phenotype; in other words, whether or not it indicates cell-type switching, is also indicated. (b) The colony-forming units of white cells expressing GFP (meaning they underwent opaque-to-white switching, blue circles) and the colony-forming units of white cells expressing mCherry (red squares) are plotted for each mouse over the course of the 17-day experiment. The raw data from this figure is available in Dataset 6. (c) Using a flowchart, the experimental setup, cell type, and potential fluorescence phenotypes for each strain are tabulated. In this case, white cells expressing mCherry and opaque cells expressing GFP were inoculated into 4 mice via oral gavage. Unlike the experiment in panels a-b, these mice were not treated with antibiotics prior to the inoculation. Fecal pellets were processed to measure white and opaque cell colonization as well as white-opaque switching. The mechanistic interpretation of each phenotype; in other words, whether or not it indicates cell-type switching, is also indicated. (d) The colony-forming units of white cells expressing GFP (meaning they underwent opaque-to-white switching, blue circles) and the colony-forming units of white cells expressing mCherry (red squares) are plotted for each mouse over the course of the seven-day experiment. The raw data from this Figure is available in Dataset 6.

The standard intestinal colonization model employed above involves a 4-day antibiotic treatment of the mice preceding the *C. albicans* inoculation. To compare white and opaque cells in the GI tract in the presence of an intact mouse microbiota, we inoculated non-antibiotic treated mice with an equal number of white and opaque cells via oral gavage (1*10^8^ cells total per mouse, n=4 mice, Figure 4c) and then tracked white and opaque cells during the infection through fecal pellets. As expected, we found that significantly fewer *C. albicans* cells colonized the GI tract when an intact microbiota was present (Figure 4d, ~100-1000 times fewer than in the standard model described above and in Figure 4a) and that the few cells that did colonize the GI tract were almost exclusively white cells that had been injected as white cells. We did not observe long-term survival of any cells that were introduced as opaque cells. These results indicate that opaque cells are out-competed by white cells in the murine gut colonization model and that this trend is more pronounced in the untreated mice than in the antibiotic-treated mice.

## Discussion

White and opaque cells are distinct, heritable cell types of *C. albicans* produced from the same genome. White cells and opaque cells differ in many characteristics and are believed to be specialized for different niches in their mammalian hosts (reviewed in Huang, 2012; Lohse and Johnson, 2009; Morschhäuser, 2010; Soll, 2014, 1992, 2004). We used variations on two murine models of *C. albicans* infection combined with isogenic fluorescent reporter strains to directly compare the fate of white and opaque cells *in vivo* and to also monitor switching between the two cell types in the host.

Many published experiments have monitored the fate of *C. albicans* white cells in murine models of infection, and our results with white cells are entirely consistent with this large body of data. While most of the experiments in the literature have utilized SC5314-derived strains, which are a/α and therefore remain locked as white cells, we used mating type **a** derivatives of SC5314, which have the ability to undergo white-opaque switching. Despite this difference, white **a** cells, in our experiments, behaved similarly to white a/α cells in the published literature. In a sense, we treated white cells as a control to understand the fate of genetically identical opaque cells. We also note that one of our observations with opaque cells, namely that opaque cells are deficient in colonizing the kidney relative to white cells, is consistent with a previous report (Kvaal et al., 1997).

To gain a better understanding of opaque cells *in vivo*, we examined four organs in addition to the kidney. Because the standard animal models of *C. albicans* infection have been optimized to study white cells (predominantly a/α cells), we also varied the dosage, timing and antibiotic treatments in these models to increase the chances of observing opaque cell behavior *in vivo*. Finally, although our *Candida* strains had been genetically altered to express different fluorescent proteins, they represent normal switching strains rather than strains locked in one form or the other. We show that during systemic infection initiated by tail-vein injection, opaque cells can disseminate into the bloodstream and colonize all five of the organs we monitored. However, the pattern of colonization across organs differs significantly from that of white cells. For example, when injected in equal numbers, white cells greatly predominated in the kidney with a substantial fraction of the white cells having initially been injected as opaque cells and undergone opaque-to-white switching. In contrast, opaque cells outnumbered white cells in the heart and spleen, perhaps our most significant observation. Roughly equivalent colonization of the two cell types was observed in the liver and brain. From this work, we have identified *in vivo* environments where opaque cells compete favorably with white cells; such environments have only rarely been identified *in vitro* (Ene et al., 2016). We anticipate that further modification of *C. albicans* infection models previously optimized for white cells will be valuable in learning about the function of opaque cells *in vivo*.

In the mouse GI model, white cells out-numbered opaque cells at every time point in every mouse. All of the cells we examined that had been introduced as opaque cells and subsequently recovered in the GI model had switched to the white cell type, further supporting the advantage of white cells for this model in wild-type cells.

Finally, we observed evidence of wide-spread opaque-to-white switching in both murine models. In the GI model, extensive opaque-to-white switching occurred early (day 1) in the colonization process. In the systemic model, we recovered white cells that had initially been injected as opaque cells from the kidney, liver, spleen, brain and heart (Figure 2, Figure S3). Although we do not know exactly when these cells switched, the results suggest that it generally occurred after opaque cells had colonized the organ in question (see results section). If this model is correct, then switching rates must differ from organ to organ ranging from little or none occurring in the heart to extensive switching in the brain and kidney. This conclusion is consistent with a number of *in vitro* studies showing that both switching rates and relative growth rates of white and opaque cells differ across media conditions (Ene et al., 2016).

Since its discovery in 1987 by Soll and colleagues, white-opaque switching has been extensively studied by numerous laboratories. However, the selective pressure that has preserved white-opaque switching across a clade of species (*C. albicans, C. dubliniensis*, and *C. tropicalis*) representing 50 million years of evolutionary time is not understood. White and opaque cells differ in their appearance, their mating ability, their recognition by macrophages, and their growth on different laboratory media. Yet, the extensive gene expression changes between the two cell types suggest that these phenotypic differences constitute only a subset of the total range of differences. In this paper, we have documented several additional phenotypic differences; specifically, differences in organ colonization. In particular, we show for the first time that opaque cells out compete white cells in two organs, the heart and spleen. It also appears that opaque-to-white switching occurs at significantly different rates in different organs. These observations strongly support the hypothesis that white and opaque cells are specialized to “match” different niches in the host. It is now possible to test this hypothesis explicitly by analyzing colonization by white and opaque cells in which different “opaque-specific” and “white-specific” genes have been deleted.

## Materials and Methods

### Plasmid construction

Plasmids for GFP or mCherry tagging of *TEF2* were constructed as follows. The last 500 bp of the *TEF2* open reading frame (ORF) (excluding the stop codon) were PCR amplified with a 5’ SphI site and a 3’ linker for either GFP or mCherry. The 500bp immediately 3’ of the *TEF2* ORF was PCR amplified between NotI and AatII sites. *C. albicans* optimized GFP (Cormack et al., 1997) was PCR amplified with a 5’ 3× Gly linker and a 3’ XhoI site. *C. albicans* optimized mCherry (Lohse and Johnson, 2016) was PCR amplified with a 5’ GRRIPGLIN linker and a 3’ XhoI site. We then merged the two *TEF2* fragments and either GFP or mCherry into one fragment during a second round of PCR. This resulted in a SphI-end of TEF2 ORF-GFP-XhoI-NotI-TEF2 3’ flank-AatII fragment and a SphI-end of *TEF2* ORF-mCherry-XhoI-NotI-TEF2 3’ flank-AatII fragment that were digested with SphI and AatII and ligated into pUC19 (NEB) to form pMBL183 and pMBL184. Following sequencing, the plasmids were then digested with XhoI and NotI and ligated with the similarly digested SAT1 flipper cassette from pSFS2A (Reuss et al., 2004) to form pMBL187 and pMBL188. Both plasmids were digested with SphI and AatII prior to transformation into *C. albicans*.

### *C. albicans* Strain construction

MLY612/613 are derived from the switching-capable AHY135 background (Hernday et al., 2013), a strain where the *HIS1* and *LEU2* markers were added back to RZY47 (Zordan et al., 2006). The untagged strains MLY537 (white) and MLY589 (opaque) are stocks of AHY135 (white) and its opaque equivalent, AHY136, respectively. AHY135 was transformed with linearized pMBL187 or pMBL188 and selected for growth on Yeast Extract Peptone Dextrose (YEPD) supplemented with 200 μg/ml nourseothricin (clonNAT, WERNER BioAgents, Jena, Germany). Insertion was verified by colony PCR against the 5’ and 3’ flanks. The *SAT1* marker was then recycled by growth in Yeast Extract Peptone (YEP) media lacking dextrose that was supplemented with 2% Maltose for 6-24 hours at 30°C. Cells were then plated on YEPD supplemented with 25 μg/ml nourseothricin and grown for 24 hours at 30°C. Loss of the *SAT1* cassette was then verified by a second round of colony PCR against the 5’ and 3’ flanks. The strain genotypes are listed in detail in Table.

MLY612/613 were grown on synthetic complete media supplemented with 2% glucose and 100 µg/ml uridine (SD+AA+Uri) at room temperature (approximately 22°C) and examined for opaque sectors, which were then isolated to make opaque glycerol stocks (MLY629/642).

### Growth Rate Data Acquisition and Assay

SD+AA+Uri overnight cultures (25°C) of GFP-tagged, mCherry-tagged, and untagged white and opaque cells were started from white or opaque colonies without visible sectors. The following morning, the overnight cultures were diluted to a density of OD_600_ = 0.01 in 100 µl of SD+AA+Uri. Growth curve assays were performed on a Tecan Spark 10M at a temperature of 25°C. Absorbance was measured for each well every 15 min for 47.25 hours (190 cycles); the plate was shaken continuously between reads. This setup allowed cells to recover from the dilution, enter log phase growth, and to eventually reach saturation. Three biological replicates (each originating from separate overnight cultures) were tested for each sample. Additionally, three technical replicates from each biological replicate were tested yielding a total of nine wells for each of the six strains tested. As a control, uninoculated (blank) wells were also included.

At each time point, the background-subtracted absorbance was calculated by subtracting the average absorbance of 24 blank wells from the absorbance of each well. For the plots in Figure S1, the natural log of the average background-subtracted absorbance for each set of three technical replicates is plotted. The specific growth rates (1/hr) in Dataset 1 were estimated for each well as the maximum slope of a smooth fit to the background-subtracted data. An R script found the maximum value for a fixed time window in every well that met the following three criteria. First, the time window had to be 4 hours long (thus, including 17 total measurements), second, the correlation coefficient (R^2^) of the fit had to be at least 0.98, third, the background-subtracted absorbance value had to be at least 0.15 for each of the measurements in the time window. The average and individual specific growth rates for each strain are tabulated in Dataset 1.

To complete our analysis, we used Welch’s *t*-test (unpaired and two tailed) to compare the growth rates of the three opaque strains with each other and of the three white strains with each other. These values are also included in Dataset 1, we note that none of the p-values for these comparisons was less than 0.1.

## Plating Assay Validation

SD+AA+Uri overnight cultures were started from two independent white (MLY613) and opaque colonies (MLY629). The overnight cultures were diluted to an OD_600_ = 0.5 in SD+AA+Uri before being loaded into a BD Accuri C6 Plus flow cytometer (BD Biosciences) to measure cell number and fluorescence. A blue (488 nm) laser was used to excite GFP and emission was detected using a 533/30 nm bandpass filter. The GFP expression of these MLY613 and MLY629 diluted cultures were measured to determine a gate where more than 99% of MLY629 (opaque-GFP) cells were above the GFP threshold and more than 99% of MLY613 (white-mCherry) cells were below the GFP threshold. The cell counts from these measurements were used to create mixtures of white-opaque populations, diluted 10-fold in SD+AA+Uri from the initial stocks, where the fraction of white cells in each population (ranging from 0-100%) were known. The fraction of white cells in these mixtures were then measured by flow cytometry (measuring the proportion of the cells that are below the previously established GFP threshold) and by plating (measuring the proportion of the resulting colonies that are white). Plating was on SD+AA+Uri at 25°C and colony morphology was scored after 3 days growth; only colonies that were completely white or opaque were counted. We note that the cells had to be diluted before plating so that only ~100 colonies would grow on each plate (making the colonies on the plate easy to score as either white or opaque). The correlation between the fraction of white cells as determined by plating and cytometry are shown in Figure S2. The linear regression of these data, calculated in Graphpad Prism 8, showed that the data fit the line *y=1.0182x-3.548* with a correlation coefficient (R^2^)=0.9949. The raw data underlying the correlation are included in Dataset 2.

### Gastrointestinal Tract Colonization Model

The procedure used was essentially as described previously (Perez et al., 2013, Rosenbach et al., 2010; White et al., 2007). In brief, female Swiss Webster mice (6 weeks old, 18–20 g) were housed two per cage and treated with antibiotics (tetracycline [1 mg/ml]), streptomycin [2 mg/ml], and gentamycin [0.1 mg/ml]) added to their drinking water throughout the experiment, beginning 4 days before inoculation. In one experiment, as indicated, mice were not treated with antibiotics prior to inoculation or for the duration of the experiment.

Prior to inoculation, *C. albicans* strains were grown for 18 h at 25°C in SD+AA+Uri liquid medium, washed twice with PBS, and counted in a hemocytometer. Mice were orally inoculated, with either 5*10^7^ or 1*10^8^ *C. albican*s cells (in a 0.1 or 0.2 ml volume) as indicated, by gavage using a feeding needle. Colonization was monitored through the collection of fecal pellets (produced at the time of collection) at various days post-inoculation and at the end of the experiment when the mice were sacrificed.

Fecal pellets were weighed and suspended in sterile PBS. The fecal homogenates were serially diluted and 100 µl was plated onto SD medium containing ampicillin (50 mg/ml) and gentamycin (15 mg/ml). The antibiotics were used to prevent the growth of any contaminating bacteria. Colony forming units were determined per gram of feces. All mice were sacrificed by CO_2_ euthanasia followed by cervical dislocation.

This study was carried out in strict accordance with the recommendations in the Guide for the Care and Use of Laboratory Animals of the National Institutes of Health. The protocol was approved by the University of California San Francisco-Institutional Animal Care and Use Committee (Protocol number AN169466-02a). All efforts were made to minimize suffering.

A summary of mouse experiments is tabulated in Table 2.

### Systemic Infection Model

Female BALB/c mice (6 weeks old, 18–20 g) were infected with the *C. albicans* inoculum via tail vein injection. Saturated *C. albicans* cultures were grown at 25°C for 18 hours in SD+AA+Uri liquid medium. Cells were washed twice with sterile saline, counted in a hemocytometer and the specified inoculum was injected via tail vein in a 0.1 ml volume. Signs of infection were monitored twice per day throughout the experimental time course. In some experiments, mice were sacrificed after 24 hours. In other experiments, mice were sacrificed upon signs of illness (including inability to ambulate, staggered gait, hypothermia or significant weight loss). All mice were sacrificed by CO_2_ euthanasia followed by cervical dislocation.

This study was carried out in strict accordance with the recommendations in the Guide for the Care and Use of Laboratory Animals of the National Institutes of Health. The protocol was approved by the Office of Ethics and Compliance: Institutional Animal Care and Use Program (Protocol number AN169466-02a). All efforts were made to minimize suffering.

A summary of mouse experiments is tabulated in Table 2.

### Organ Processing and Visualization

After dissection, organs were harvested, flash frozen, and placed in a −80° C freezer. Organs were thawed on ice, homogenized in 1 ml PBS, serially diluted in PBS, and plated onto SD medium containing ampicillin (50 mg/ml) and gentamycin (15 mg/ml). The antibiotics were used to prevent the growth of any contaminating bacteria. Colonies were visualized for fluorescence using a Leica stereomicroscope. Colony forming units were determined per gram of organ.

### Animal Model Data Analysis

The colony forming units per organ are tabulated in Datasets 3-6. The raw data were normalized to the inoculum such that the inoculum was always 50:50 (this is why some of the numbers are not whole numbers). These normalized colony counts were plotted in all figures and were used to determine whether or not there were statistical differences between different distributions. First, we tested whether or not there were statistical differences between cells that were injected as white and cells that were injected as opaque. In other words, irrespective of whether the cell types switched to another cell-type after they were injected, we tested whether or not there was a difference in their initial colonization. These two distributions (normalized colony counts of cell that were injected white and normalized colony counts of cells that were injected opaque) were compared with a non-parametric test, the Wilcoxon matched-pairs signed rank test, to generate p-values and determine significance. Next, we tested whether or not there were statistical differences between the stability of the cell types. In other words, we compared two distributions (normalized colony counts of cells that were injected white and remained white and normalized colony counts of cells that were injected opaque and remained opaque) using the same non-parametric statistical test (Wilcoxon matched-pairs signed rank test) used above. The statistical analysis was conducted independently for each experiment and for each organ with GraphPad Prism 7. The raw data, normalized data and p-values are reported in Datasets 3-6.

## Acknowledgements

We thank the Preclinical Therapeutic Core Facility, and especially Donghui Wang for help with tail-vein injections and oral gavage, respectively. We thank Naomi Ziv for the R script that was used to analyze the plate reader data in Figure S1 and Dataset 1. We thank the Li Lab for use of the Leica stereomicroscope and members of the Johnson lab for discussion.

## Supporting Information Captions

**Figure S1:**
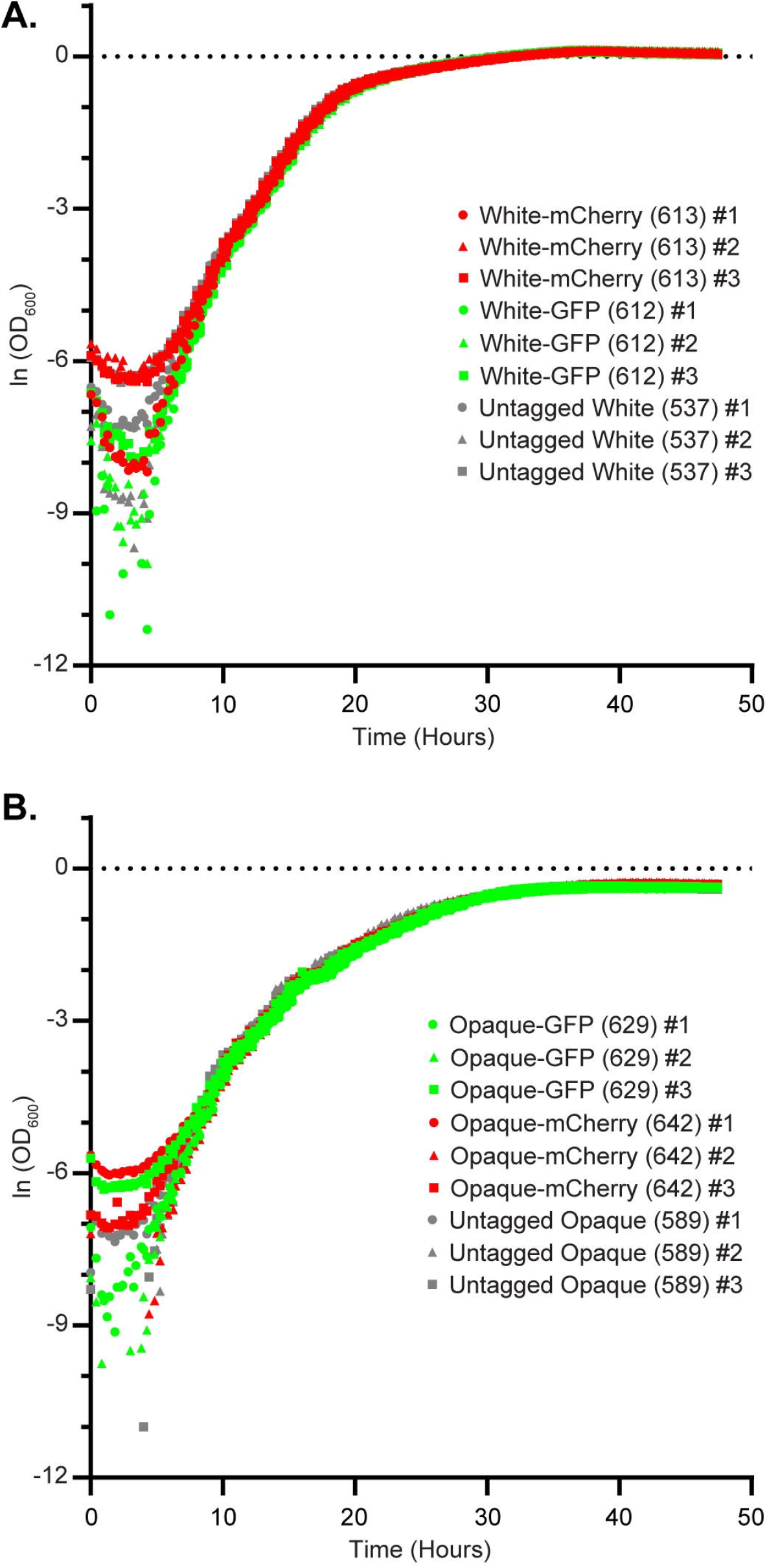
Fluorescently tagging white and opaque cells does not impact their growth rate. (a) White cells without a fluorescent tag (grey), white cells expressing GFP (green), and white cells expressing mCherry (red) were grown in SD+AA+Uri at 25°C. (b) Opaque cells without a fluorescent tag (grey), opaque cells expressing GFP (green), and opaque cells expressing mCherry (red) were grown in SD+AA+Uri at 25°C. In both panels the natural logarithm of the background subtracted absorbance at 600 nm, averaged from three technical replicates, is plotted as a function of time for the three biological replicates of each strain.

**Figure S2:**
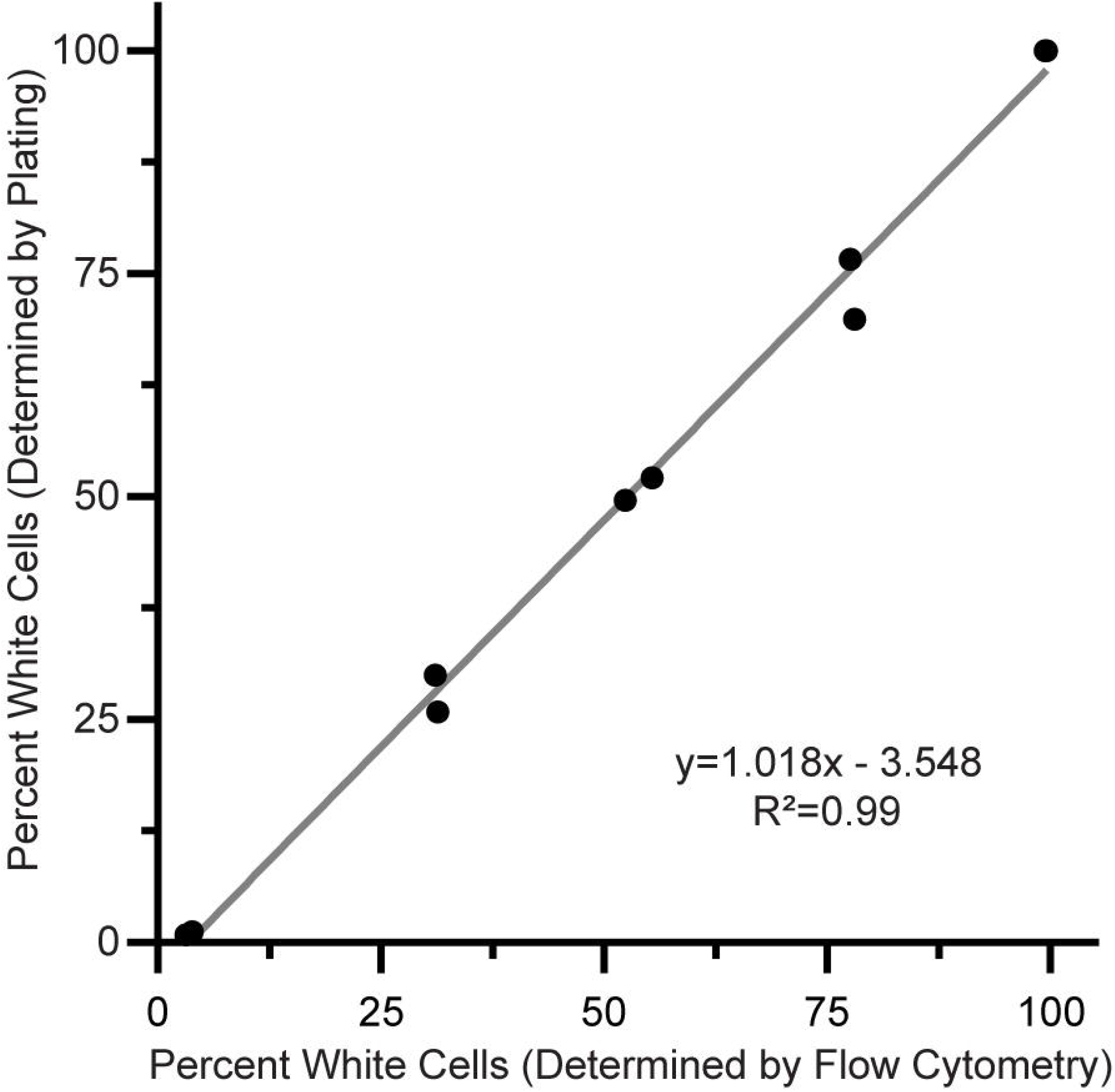
Flow cytometry shows that our plating assay accurately captures the distribution of co-cultured white and opaque cells. Opaque-GFP and white-mCherry cultures were independently grown. The cell density and fraction of the population expressing GFP (that is, opaque cells) or not expressing GFP (that is, white cells) were independently measured using a flow cytometer. The white and opaque cultures were then mixed at different ratios and the fraction of the population not expressing GFP (that is, white cells) was determined for each mixture using a flow cytometer. The mixtures were then plated and subsequently scored for colony phenotype. The proportion of the culture that did not express GFP (that is, white cells) was compared to the proportion of white colonies. The linear regression is indicated in grey.

**Figure S3.**
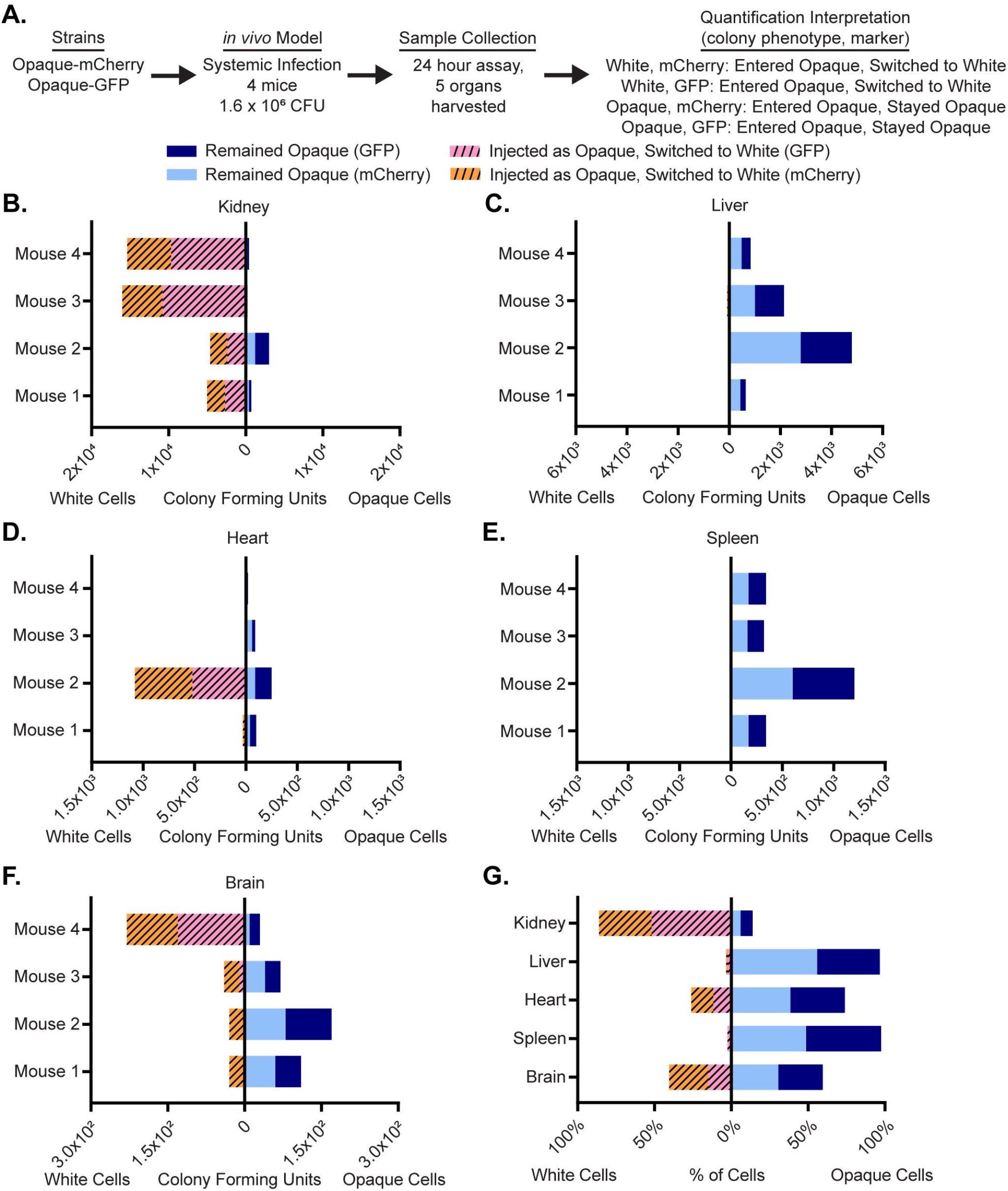
This data is related to Figure 2: Opaque cells can colonize organs without being co-injected with white cells. (a) Using a flowchart, the experimental setup, cell type, and potential fluorescence phenotypes for each strain are tabulated. In this case, opaque cells expressing mCherry and opaque cells expressing GFP were co-injected into the tail-veins of 4 mice. Five organs, the kidney, liver, heart, spleen and brain, were processed to measure white and opaque cell colonization as well as white-opaque switching. The mechanistic interpretation of each phenotype; in other words, whether or not it indicates cell-type switching, is also indicated. The colony-forming units of cells that remained opaque (i.e. opaque cells expressing mCherry (light blue) or GFP (blue)) and of cells that switched from opaque-to-white (i.e. white cells expressing GFP (dashed pink) or mCherry (dashed orange)) are plotted for each mouse for the (b) kidney, (c) liver, (d) heart, (e) spleen and (f) brain. The left side of each horizontal bar graph refers to cells that were white at the end of the experiment while the right side of each horizontal bar graph refers to cells that were opaque at the end of the experiment. (g) The mean percentage of total cells that remained opaque (i.e. opaque cells expressing mCherry (light blue) or GFP (blue)) or that switched from opaque-to-white (i.e. white cells expressing GFP (dashed pink) or mCherry (dashed orange)) are plotted per organ as a horizontal bar graph. The left side of the horizontal bar graph refers to cells that were white at the end of the experiment while the right side of the horizontal bar graph refers to cells that were opaque at the end of the experiment. The raw data for this experiment is available in Dataset 4.

**Figure S4.**
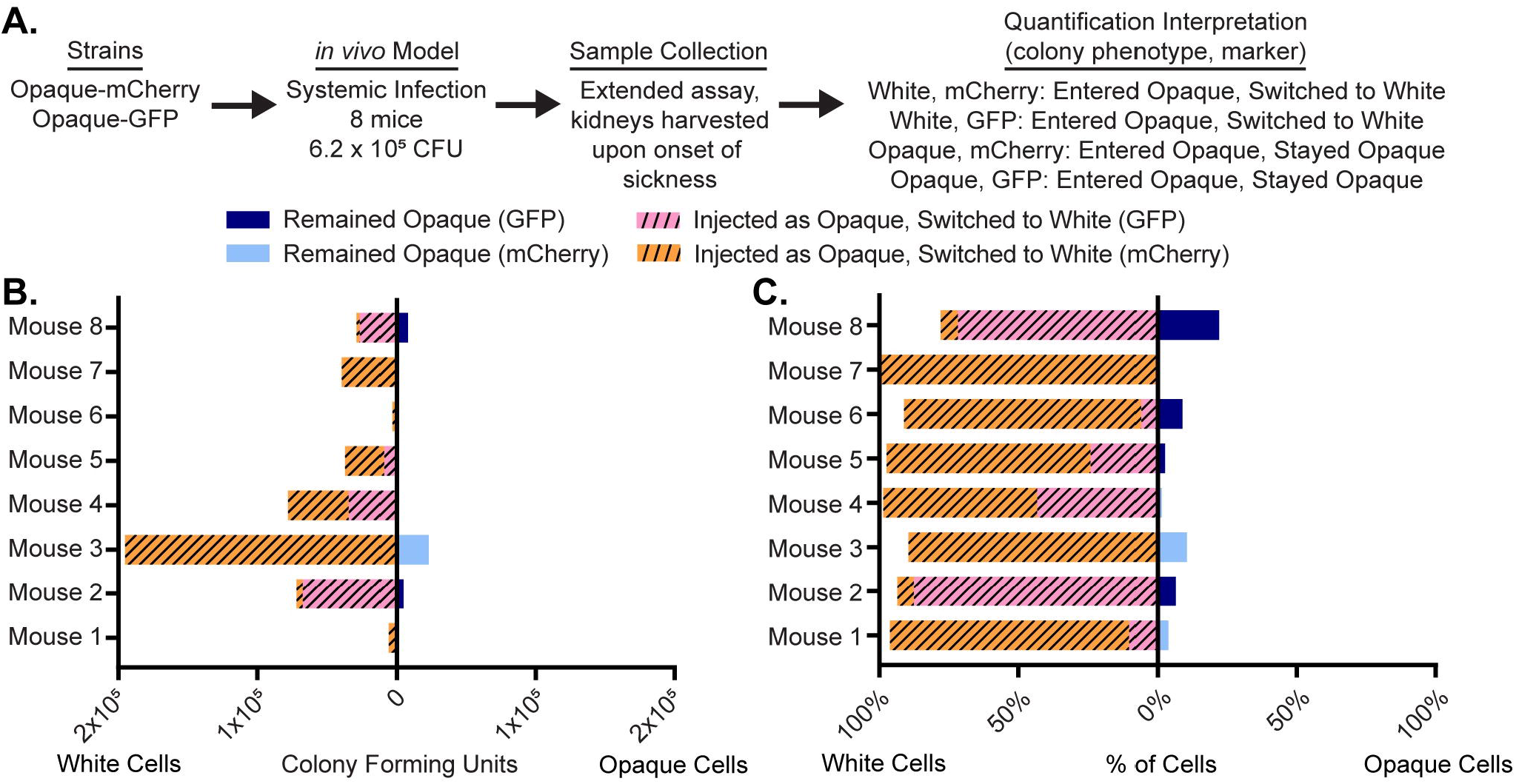
This figure is related to Figure 3: Opaque cells switch to white cells in the kidney. (a) Using a flowchart, the experimental setup, cell type, and potential fluorescence phenotypes for each strain are tabulated. In this case, opaque cells expressing mCherry and opaque cells expressing GFP were co-injected into the tail-veins of 8 mice. Upon the onset of illness, the kidney was processed to measure opaque cell colonization as well as opaque-to-white switching. The mechanistic interpretation of each phenotype; in other words, whether or not it indicates cell-type switching, is also indicated. (b) The colony-forming units of cells that remained opaque (i.e. opaque cells expressing mCherry (light blue) or GFP (blue)) and of cells that switched from opaque-to-white (i.e. white cells expressing GFP (dashed pink) or mCherry (dashed orange)) are plotted per mouse as a horizontal bar graph. The left side of the horizontal bar graph refers to cells that were white at the end of the experiment while the right side of the horizontal bar graph refers to cells that were opaque at the end of the experiment. (c) The percentage of total cells that remained opaque (i.e. opaque cells expressing mCherry (light blue) or GFP (blue)) or that switched from opaque-to-white (i.e. white cells expressing GFP (dashed pink) or mCherry (dashed orange)) are plotted for each mouse as a horizontal bar graph. The left side of the horizontal bar graph refers to cells that were white at the end of the experiment while the right side of the horizontal bar graph refers to cells that were opaque at the end of the experiment. The raw data for this experiment is available in Dataset 5.

**Figure S5.**
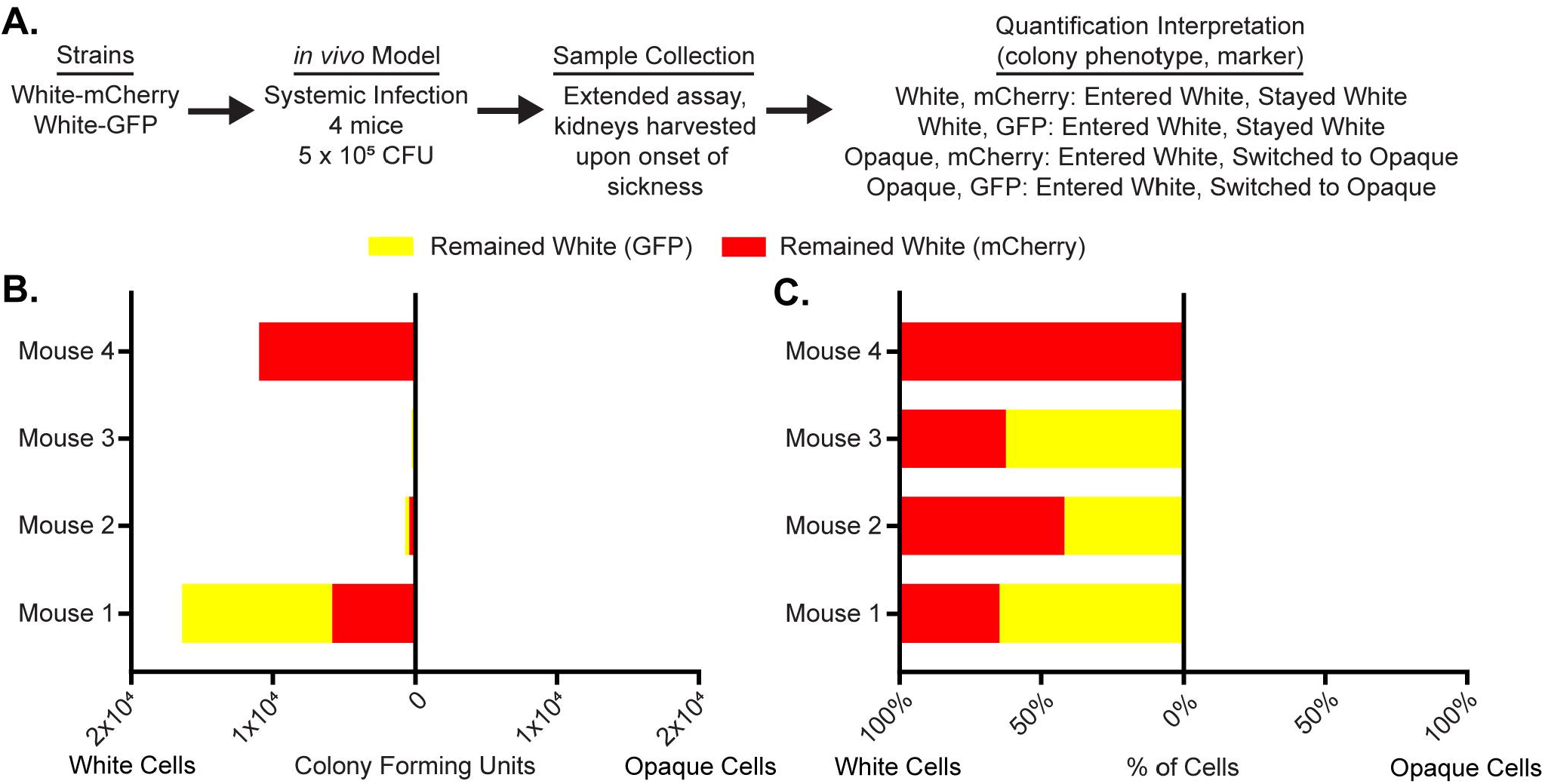
related to Figure 3: White cells are stable in the kidney. (a) Using a flowchart, the experimental setup, cell type, and potential fluorescence phenotypes for each strain are tabulated. In this case, white cells expressing mCherry and white cells expressing GFP were co-injected into the tail-veins of 4 mice. Upon the onset of illness, the kidney was processed to measure white cell colonization as well as white-opaque switching. The mechanistic interpretation of each phenotype; in other words, whether or not it indicates cell-type switching, is also indicated. (b) The colony-forming units of white cells expressing GFP (yellow) and the colony-forming units of white cells expressing mCherry (red) are plotted per mouse as a horizontal bar graph. The left side of the horizontal bar graph refers to cells that were white at the end of the experiment while the right side of the horizontal bar graph refers to cells that were opaque at the end of the experiment. (c) The percentage of total cells that remained white (i.e. white cells expressing mCherry (red) or GFP (yellow)) are plotted for each mouse as a horizontal bar graph. The left side of the horizontal bar graph refers to cells that were white at the end of the experiment while the right side of the horizontal bar graph refers to cells that were opaque at the end of the experiment. The raw data for this experiment is available in Dataset 5.

**Table 1:**
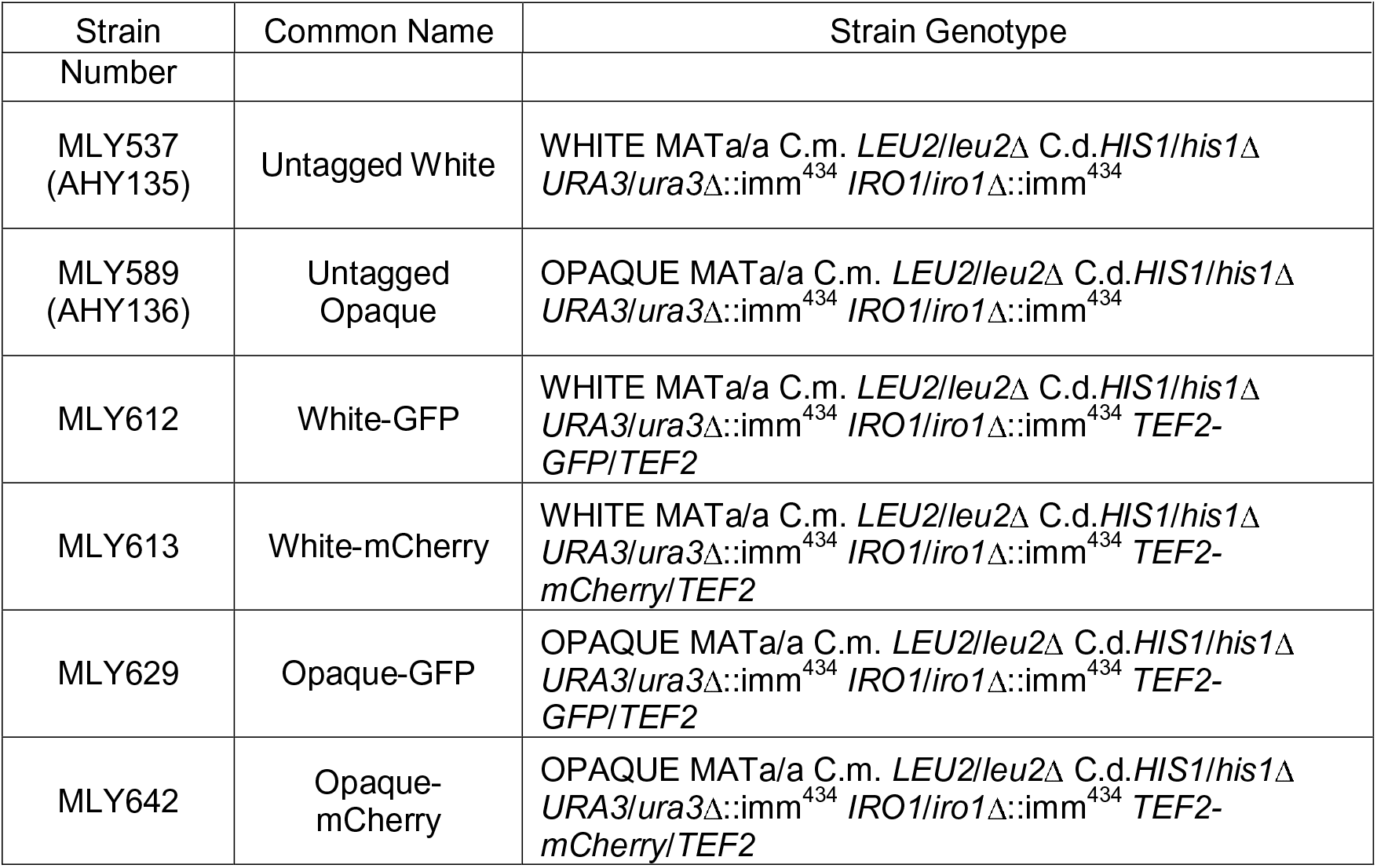
*C. albicans* strains used in this study

**Table 2:** Summary of *in vivo* experiments conducted in this work

| Inoculum Strain 1 | Inoculum Strain 2 | Infection Type (Figure #) | Total Inoculum | Duration | Number of Mice | Samples Harvested |
| --- | --- | --- | --- | --- | --- | --- |
| White-GFP | White-mCherry | Systemic (Figure 1) | $2.2 \times 10^6$ | 24 hours | 4 | 5 Organs |
| White-mCherry | Opaque-GFP | Systemic (Figure 2) | $8.0 \times 10^5$,<br>$1.6 \times 10^6$ | 24 hours | 8 (4 Each Inoculum) | 5 Organs |
| Opaque-GFP | Opaque-mCherry | Systemic (Figure S3) | $1.6 \times 10^6$ | 24 hours | 4 | 5 Organs |
| White-mCherry | Opaque-GFP | Systemic (Figure 3) | $5.0 \times 10^5$ | Variable (Onset of Sickness) | 7 | Kidneys |
| Opaque-GFP | Opaque-mCherry | Systemic (Figure S4) | $6.2 \times 10^5$ | Variable (Onset of Sickness) | 8 | Kidneys |
| White-GFP | White-mCherry | Systemic (Figure S5) | $5.0 \times 10^5$ | Variable (Onset of Sickness) | 4 | Kidneys |
| White-mCherry | Opaque-GFP | Intestinal Colonization (Figure 4B) | $5.0 \times 10^7$ | 17 days | 8 | Fecal Pellets |
| White-mCherry | Opaque-GFP | Intestinal Colonization, No Antibiotic (Figure 4D) | $1.0 \times 10^8$ | 7 days | 4 | Fecal Pellets |

**Dataset 1: Raw data and statistical analysis of data plotted in Figure S1**.

**Dataset 2: Raw data for Figure S2.**

**Dataset 3: Raw data, normalized data, and statistical analysis of data plotted in Figure 1.**

**Dataset 4: Raw data, normalized data, and statistical analysis of data plotted in Figures 2 and S3.**

**Dataset 5: Raw data, normalized data, and statistical analysis of data plotted in Figures 3, S4 and S5.**

**Dataset 6: Raw data, normalized data, and statistical analysis of data plotted in Figure 4.**

